# Collapse of Interictal Suppressive Networks Permits Seizure Spread

**DOI:** 10.1101/2024.12.04.626580

**Authors:** Ghassan S. Makhoul, Derek J. Doss, Graham W. Johnson, Price C. Withers, Addison C. Cavender, Bruno Hidalgo Monroy Lerma, Anas Reda, Camden E. Bibro, Emily Liao, Hakmook Kang, Benoit M. Dawant, Shilpa B. Reddy, Angela N. Crudele, Christos Constantinidis, Shawniqua Williams Roberson, Sarah K. Bick, Victoria L. Morgan, Dario J. Englot

**Author notes:** Correspondence to: Dario J. Englot Vanderbilt University, 1500 21st Avenue South, VAV 4340, Nashville, Tennessee 37212.

## Abstract

How do networks in the brain limit seizure activity? In the Interictal Suppression Hypothesis (ISH), we recently postulated that high inward connectivity to seizure onset zones (SOZs) from non-involved zones (NIZs) is a sign of broader network suppression at rest. If broad networks appear to be responsible for interictal SOZ suppression, what changes during seizure initiation, spread, and termination? For patients with drug resistant epilepsy, intracranial monitoring offers a view into the electrographic networks which organize around and in response to the SOZ. In this manuscript, we investigate network dynamics in the peri-ictal periods to assess possible mechanisms of seizure suppression and the consequences of this suppression being overwhelmed. Peri-ictal network dynamics were derived from stereo electroencephalography (SEEG) recordings from 75 patients with drug-resistant epilepsy undergoing pre-surgical evaluation at Vanderbilt University Medical Center. We computed directed connectivity from 5-second windows in the periods between, immediately before, during, and after seizures. After aligning all network connectivity matrices between seizures and patients, we calculated net connectivity changes from the SOZ, propagative zone (PZ), and NIZ. Across all seizure types, we observed two distinct phases as seizures initiated and evolved: a large rapid increase in directed communication towards the SOZ from NIZ followed by a collapse in network connectivity. During this first phase, SOZs could be distinguished from all other regions (One-Way ANOVA, p=8.32×10^−19^ - 2.22×10^−7^, lower range to upper range of p-values). In the second phase and post-ictal period, SOZ inward connectivity decreased yet remained distinct (One-Way ANOVA, p= 2.58×10^−10^-1.66×10^−2^). Furthermore, NIZs appeared to drive the increase in inward SOZ connectivity while global connectivity between NIZs concordantly decreased. Stratifying by seizure subtype, we found that consciousness-impairing seizures show loss of inward connectivity from the NIZ earlier than conscious sparing seizures (one-way ANOVA, p<0.01 after false discovery correction). Tracking network reorganization against a surrogate for seizure involvement highlighted a possible antagonism between seizure propagation to the NIZ and the NIZ’s ability to maintain high connectivity to the SOZ. Finally, we found that inclusion of peri-ictal connectivity improved SOZ classification accuracy from previous models to a combined area under the curve of 93%. Overall, NIZs appear to actively respond to seizure onset and increase inhibitory signaling towards the SOZ, possibly in an attempt to thwart seizure activity. This inhibition appears to be insufficient to prevent seizure onset, and furthermore, loss of normal communication in the rest of the brain between NIZs may contribute to loss of consciousness during larger seizures. Dynamic connectivity patterns uncovered in this work may: i) allow more accurate delineation of surgical targets in focal epilepsy, ii) reveal why inward suppression of SOZs interictally may nonetheless be insufficient to prevent all seizures, and iii) provide insight into mechanisms of loss of consciousness during certain seizures.

## Introduction

Epilepsy is a common neurological disorder affecting 1% of the global population with drug resistant epilepsy (DRE) comprising 40% of patients.^1-3^ For these patients, surgical intervention can be potentially curative, however approximately 30-40% continue to live with recurrent seizures even after surgical intervention.^4^ To improve the rate of curative epilepsy surgery, extensive pre-operative investigations are performed to localize the area thought to be generating seizures, termed the seizure onset zone (SOZ), as successful resection of the SOZ may lead to seizure freedom. While this approach works for some patients, for others SOZ resection alone is not enough to produce seizure freedom.^5-7^ Thus, research has focused on identifying the seizure network and disrupting it.

For over 20 years, epilepsy has been described as a network pathology, which departs from the idea that seizures are the result of a local diseased population of neurons.^8^ Instead, the network theory posits that epilepsy remodels brain networks into pathologic states, and that seizures are an expression of this network pathology. Furthermore, the network theory orients investigation toward studying global disruption within epilepsy: white matter loss, functional changes, neurocognitive deficits, etc.^9-12^ Researchers have developed electrographic descriptions of epilepsy networks by constructing functional connectivity networks from stereo electroencephalography (SEEG) recordings from patients with DRE undergoing intracranial monitoring for pre-surgical evaluation.^13-15^ Prior work has shown that SOZs exhibit high net inward connectivity from the broader network, specifically the non-involved zones (NIZ).^16-19^ In the Interictal Suppression Hypothesis (ISH), we postulated that this high net inward connectivity is a signature of SOZ suppression by the broader network that helps explain with patients with focal epilepsy are not constantly having seizures.^20, 21^

These interictal studies would then imply that reconfigurations between the SOZ and the broader network precipitate seizures. Prior work investigating interictal to ictal network transitions has found that the transition into the ictal state is not solely a function of imbalanced excitation-inhibition, but instead may be the result of a failed feedback mechanism or decoupling to the SOZ.^22, 23^ Additional work in consciousness-impairing seizure dynamics suggests that peri-ictal networks with higher thalamocortical coupling yield more severe loss of conscioussness.^24^ Thus given that our prior work in the ISH characterized the SOZ as a zone of high feedback inhibition, we hypothesized that these peri-ictal findings may be the result of a failure in interictal suppression. Specifically, if the ISH can explain the maintenance of network homeostasis, then does this imply that diminishing interictal suppression creates a permissive network for seizures? Furthermore, would this same directed network play any role in seizure dynamics? We hypothesized that seizure onset would be coincident to a change in interictal suppression, and that seizure propagation may be explained by the same suppressive signal being overwhelmed. We expected that the degree of feedback degeneration to the SOZ would correspond to seizure spread. Conversely, seizure spread may be mitigated by networks which can successfully maintain high feedback to the SOZ throughout the ictal period. As such, we propose that SOZ suppression may be a general feature of brain networks and thus the ISH merits an extension into the peri-ictal period as a general inhibitory network. Outside of evaluating this generalization of the ISH, studying peri-ictal dynamics can be a valuable source for improving SOZ localization and thus guiding surgical intervention.^25^

To evaluate our hypothesis, we sought to assess peri-ictal dynamics in the following manners: (1) We aimed to relate changes in network suppression to transitions into and out of the seizure state. (2) We aimed to describe seizure specific network dynamics by stratifying focal aware seizures (FAS), focal impaired awareness seizures (FIAS), and focal to bilateral tonic-clonic (FBTC) seizures. (3) We then sought to assess whether the SOZ inward connectivity was a potential mechanism to constrain seizure propagation. (4) Finally, we evaluated the direct translation of these metrics into clinical utility. This work may assist in further localization of the SOZ and may increase our understanding of the mechanisms of seizure initiation, spread, and termination. Finally, if we can find areas of the network which antagonize seizure spread, then we may be able to shift from resecting problematic areas to assisting adaptive network configurations through neuromodulation.^26^

## Materials and Methods

### Participants, Data Acquisition and Pre-processing

We analyzed SEEG data collected from 75 patients with DRE who were evaluated at the Vanderbilt University Medical Center. Their demographic and clinical data are outlined in **Table I**. This study was approved by Vanderbilt’s Institutional Review Board and all patients provided informed consent. The treating neurosurgeons developed trajectories for electrode placement (Ad-Tech, Oak Creek, WI or PMT Corporation, Chanhassen, MN) using waypoint software (FHC, Bowdoin, ME). Bipolar pairs in gray matter were localized with CRAnial Vault Explorer software (CRAVE; Vanderbilt University, Nashville, TN), as in prior work.^16, 20, 27^

**Table 1.** Subject demographics and clinical Information. Data are N (%) for count variables and mean ± SD for continuous variables. ATL = anterior temporal lobectomy; DBS = deep brain stimulation of bilateral anterior thalamic nuclei; FBTC = focal to bilateral tonic-clonic seizures; mTLE = mesial temporal lobe epilepsy; MTS = mesial temporal sclerosis; PVNH = Periventricular Nodular Heterotopia; RNS = responsive neurostimulation; SAH = selective amygdalohippocampectomy; TLE = temporal lobe epilepsy. N = 75 patients.

|  |  |
| --- | --- |
| Sex, female | 43 (57.3%) |
| Age, years | 34.7 ± 12.3 |
| Epilepsy durations, years |  |
| FBTC seizures, yes (n, %) | 58 (77.3%) |
| MTS, yes | 19 (25.3%) |
| **Clinically Presumed epilepsy subtype (n, %)** |  |
| Unilateral mTLE | 21 (28%) |
| Bilateral mTLE | 9 (12%) |
| Unilateral lateral TLE | 17 (26%) |
| Unilateral frontal lobe epilepsy | 6 (8%) |
| Unilateral parietal lobe epilepsy | 1 (1.3%) |
| Multiple presumed foci (not mTLE) | 15 (20%) |
| Insular | 2 (2.6%) |
| PVNH/Cavernous Malformation | 2 (2.6%) |
| Encephalomalacia | 2 (2.6%) |
| Unknown | 3 (4%) |
| **Surgery type (n, %)** |  |
| Resective SAH | 14 (18.7%) |
| ATL | 10 (13.3%) |
| Other resection | 11 (14.7%) |
| Laser SAH | 4 (5.3%) |
| Neurostimulation (DBS/RNS) | 26 (34.7%) |
| None | 10 (13.3%) |
| **SEEG node designation** | 84.9 ± 26.1 |
| SOZ | 15.7 ± 14.2 |
| PZ | 27.3 ± 22.6 |
| NIZ | 189.2 ± 75.0 |
| **Seizure Events** |  |
| FAS | 123 (18.4%) |
| FIAS | 149 (22.3%) |
| FBTCS | 82 (12.3%) |
| Focal Unknown Awareness | 314 (47%) |
| Total | 668 (100%) |

All patients were monitored in the Vanderbilt epilepsy monitoring unit (EMU) with standard video-SEEG protocols. Epileptologists designated channels as belonging to the SOZ, PZ, or NIZ using the following criteria: Channels which capture the earliest electrographic activity of a seizure were deemed the SOZ. Channels which display seizure activity within 10 seconds of initiation, but not at seizure onset were deemed propagation zones (PZ). The remaining channels were designated as NIZs. Importantly, these designations are specific for each seizure, allowing for variation in SOZ and PZ at a per seizure level. This resulted in a total of 668 seizures analyzed.

For each seizure we collected ten minutes of recordings prior to seizure onset, the entire ictal period, and ten minutes after seizure offset. The pre-ictal period was defined as the minute prior to seizure onset, with the remaining 9 minutes of pre-seizure data defined as the interictal period. We divided the seizure period into two periods: an early ictal and late ictal period. The early ictal period was defined as the first second of seizure onset up until the timestamp marking half of seizure duration. We defined the late ictal period from this halfway point through seizure termination. We analyzed seizures with at least thirty seconds of data. Finally, we analyzed the first minute after electrographic offset as the post-ictal period to balance against the one minute of pre-ictal data. Each SEEG channel was re-referenced as a bipolar montage. We then filtered SEEG channels using MATLAB’s filtfilt function (MathWorks Inc. Natick, MA, USA). We filtered the data with a 1-59, 61-119, 121-159Hz bandpass with a 5^th^ order Butterworth filter to remove direct current shifts. The resulting bipolar pairs were then used for connectivity calculations. From here on, we will refer to these bipolar pairs as nodes.

### Peri-Ictal Connectivity Calculations

Using 5 seconds of time series data from all SEEG channels in gray matter (**Fig. 1A**), we computed directed connectivity using partial directed coherence (PDC) across 5 canonical frequency bands, delta (1-4 Hz), theta (4-8 Hz), alpha (8-12 Hz), beta (12-30 Hz), low gamma low (30-80 Hz), and high gamma (80-150 Hz). As in our previous work, these connectivity metrics were largely stable throughout low and high frequency bands, and thus we focussed analysis to the alpha band.^20, 21, 28^ We used a 1 second stride to generate dynamic connectivity for all peri-ictal periods **(Fig. 1B, C)** As in previous work, we defined inward connectivity as the average inward PDC toward a node in each window. Outward connectivity was defined as the average outward PDC from a node in each window. Net connectivity was calculated by subtracting outward from inward connectivity. Each connectivity matrix was then aggregated by region designation (**Fig 1D**), then aligned across patients to examine changes in seizure onset.

**Fig 1.**
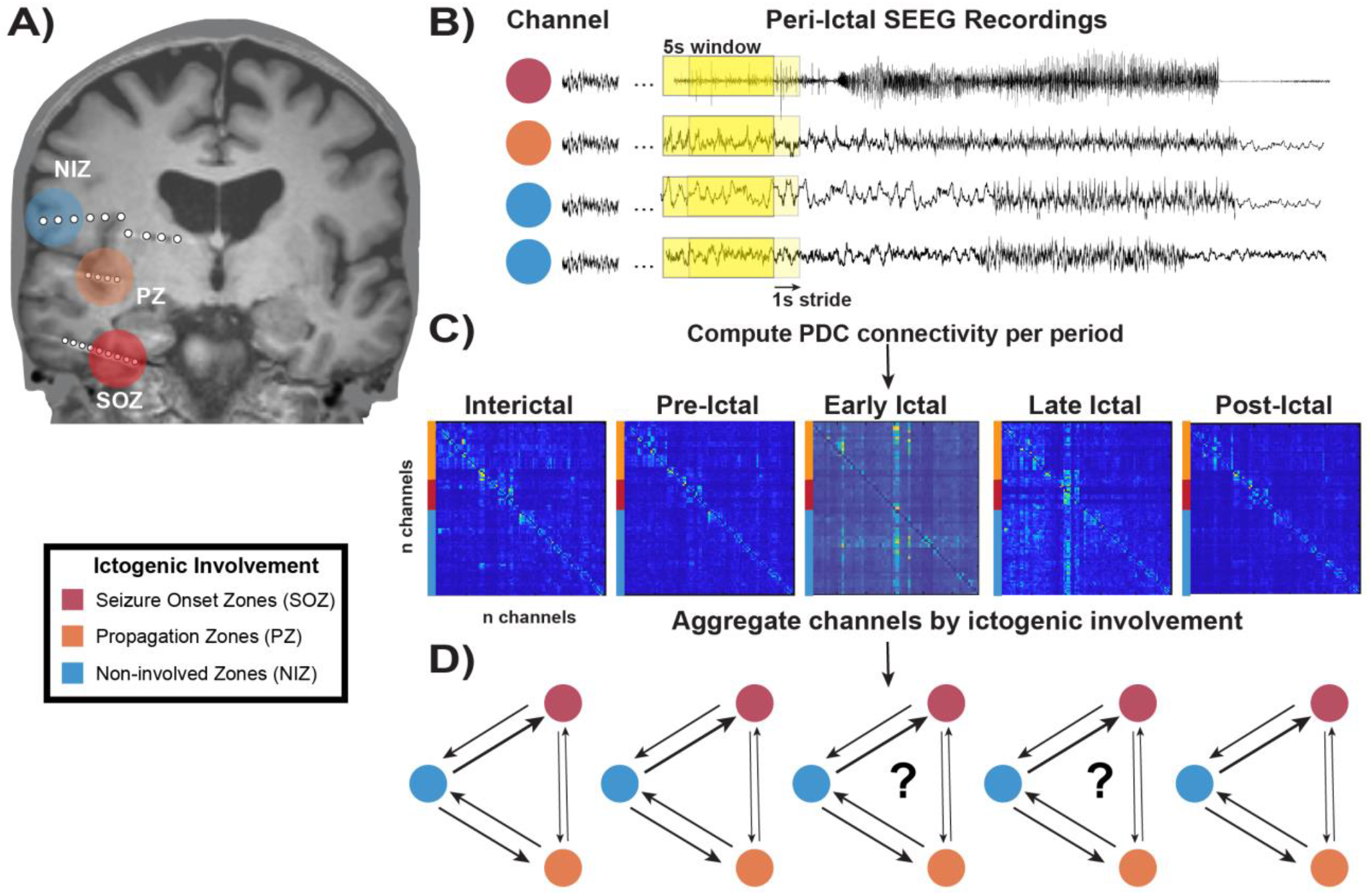
Data collection and peri-ictal network construction schematic. **A)** 75 patients were implanted with stereo electroencephalography (SEEG) leads and monitored in the epilepsy monitoring unit to capture seizure activity. **B)** Epileptologists designate channels which capture seizure initiation as seizure onset zones (SOZ), propagative zones (PZ) as channels with ictal spread within 10s of onset, and all other nodes as non-involved zones (NIZ) **C)** A 5-second sliding window with 1-second stride was used to calculate partial directed coherence connectivity matrices for all periods of interest: interictal, pre-ictal, early ictal, late ictal and post-ictal. **D**) Using epileptologist designations, dynamics between the three regions of interest are aggregated and summarized for each period of interest.

### Peri-Ictal Network Aggregation

Using timestamps from epileptologist readings, we aligned seizure onset across patients. The connectivity matrix which captures the first second of seizure activity was deemed seizure onset. We analyzed all peri-ictal data where the seizure type was known through clinical testing, and the electrographic seizure lasted at least 30 seconds. To highlight transitions into and out of the ictal states, we preserved the first and last 10 seconds of all seizure periods for all patients. To handle differences in seizure length, we aggregated the middle period of the seizure so that all patient’s had 10 seconds worth of network dynamics. For example, if a seizure lasted for 50 seconds, we would preserve the first 10 seconds worth of connectivity matrices and the last 10 seconds of connectivity matrices. For the middle period, we would average three seconds worth of connectivity matrices at a time to generate a 10 second aggregate of the middle window. For each period, we calculated inward, outward, and net connectivity of the SOZ, PZ, and NIZ and plotted these metrics over the whole group for each window with 95% confidence intervals. We compared regional connectivity profiles within each window to assess if the same motif we calculated in prior work on the ISH was preserved across periods and could continue to distinguish the SOZ, PZ, and NIZ throughout the peri-ictal period. We used a one-way ANOVA with subsequent paired t-test with Holm Bonferroni multiple comparison corrections to assess significance with alpha defined as .05.

### Seizure Subtype Analysis

For each seizure, ictal awareness testing was performed as soon as the medical staff were alerted to seizure activity as in prior work.^29^ Ictal video was reviewed to verify awareness testing and definitively label seizures as FAS, FIAS, or FBTCS. When awareness testing was equivocal or ictal video was obstructed, seizures were not given a definitive label. These seizures were excluded from analysis. To examine differences in network dynamics among seizure subtypes, we calculated net connectivity to the SOZ for FAS, FIAS, and FBTCS and aligned these connectivity matrices in a similar manner as before. Aggregating over each seizure per subject, per window, we used a one-way ANOVA test to compare network trajectories to determine if there were periods of significant divergence. We used false discovery rate to correct for the multiple windows compared. To analyze if seizure length across seizure types contributed to connectivity profiles, we calculated and compared seizure lengths across all three seizure types.

Finally, we computed an aggregate of net (inward – outward) SOZ connectivity during the ictal period for all three seizure types.

### Describing the Regions Driving Seizure Dynamics

To track changes in ictal communication patterns, we examined the NIZ’s connectivity to the SOZ, PZ and itself during the pre-ictal, ictal, and post-ictal periods. We z-scored all channels against their own inter-ictal edge weights then summarized, aggregated, and averaged within each designation (SOZ, PZ, NIZ). Per channel z-scoring allowed us to form a uniform baseline between all regions in the interictal period to compare ictal changes in communication dynamics against the interictal period and control for the already high communication from the NIZ to the SOZ as observed in the ISH. Aggregating then averaging over region designation was also designed to control for higher sampling of the NIZ compared to the PZ and SOZ. We then plotted average regional communication from the NIZ to all three regions over the ictal period

### Examining Network Dynamics Against Seizure Involvement

Next, we sought to relate NIZ recruitment against changes in network suppression. Beta power has been shown to act as a surrogate marker of seizure activity, specifically seizure spread.^30, 31^ Thus, we analyzed NIZ communication dynamics against changes in peri-ictal beta power of the NIZ z-scored against the interictal baseline beta power. We calculated bipole channel level beta-power using MATLAB’s pwelch implementation over the same 5 second windows then binned power into the canonical frequency bands with beta defined as 13-30Hz. We aligned changes in beta with network dynamics using a similar window labelling schema as described above. We computed the point of inflection using second order differencing of instantaneous beta power to generate an estimate of the period of maximum seizure propagation. We compared transition to rapid beta propagation with network trajectories. Finally, to quantify the interaction between spreading beta power and NIZ communication dynamics we used a repeated measure correlation regression between the z-scored changes in NIZ communication to the SOZ and the absolute value of the z-scored beta power.^32^

### Classifying the SOZ with Peri-Ictal Connectivity in a Patient Specific Manner

We aimed to assess the clinical utility of analyzing network dynamics by constructing a machine learning model, the support vector classifier (SVC), for each patient’s connectivity matrices. We trained one model on data from patients with Engel I outcomes (n=23) and another model on the full dataset, including patients with Engel II-IV (n=14) outcomes, patients who received neuromodulation (n= 26), and those without outcome scores or those who did not undergo surgery (n=12). For each patient, we aggregated net node-level connectivity (inward – outward) for all periods of interest. We then used 4-fold nested cross validation and class rebalancing to optimize each patient’s SVC, similar to our previous work.^20^ During each inner fold of cross validation, a grid search over parameters was performed to optimize each model. The highest performing model was then trained on all the cross-validation data and assessed against the held-out test set. We then aggregated over all models to generate average performance over the whole group. We assessed each model’s receiver operating characteristic (ROC) curve to determine its accuracy in successfully classify the SOZ vs. all other regions.

## Results

### Peri-Ictal Connectivity Highlights Two Seizure Phases

In the interictal phase, the SOZ and PZ demonstrated high inward connectivity, recapitulating prior findings in the ISH (**Fig. 2 A, C**).^20^ Interestingly, this pattern of connectivity is stable until seizure onset, even remaining stable directly before ictal onset. During the early ictal phase there is a stark increase in inward connectivity toward the SOZ and PZ while inward connectivity decreases for the NIZ (**Fig. 2 A, B**). In the late ictal phase, inward and outward connectivity precipitously decreases across all three regions (**Fig. 2 A-C**). Despite this decrease in inward connectivity, net connectivity (inward – outward) for SOZ and PZ continues to remain biased towards inward connectivity. All three regions demonstrate significant differences in net connectivity for all periods (**Fig. 2C**, One-Way ANOVA, p=8.32×10^−19^ - 2.22×10^−07^, lower range to upper range of p-values) and demonstrate pairwise differences for all three regions on two sample t-tests for the interictal, pre-ictal, early ictal periods (p=4.68×10^−18^ – 2.2×10^−4^). During the late ictal period the PZ and SOZ exhibit significantly different connectivity from the NIZ (p=2.62×10^−7^, 5.34×10^−12^ respectively). However, the SOZ and PZ cannot be distinguished from each other during this period (p=5.7×10^−2^, multiple comparison cutoff 1.25×10^−2^). During the post-ictal period, only SOZ net connectivity is significantly different from the NIZ (p=8×10^−3^) and PZ (p=8×10^−3^). The PZ and NIZ demonstrate no significant differences between each other. These findings suggest a bi-phasic network profile in the ictal period: in phase 1 net inward connectivity to the SOZ dramatically rises. In phase 2, global connectivity precipitously collapses with a marked undershoot of net inward connectivity to the SOZ which carries through the post-ictal period.

**Fig. 2:**
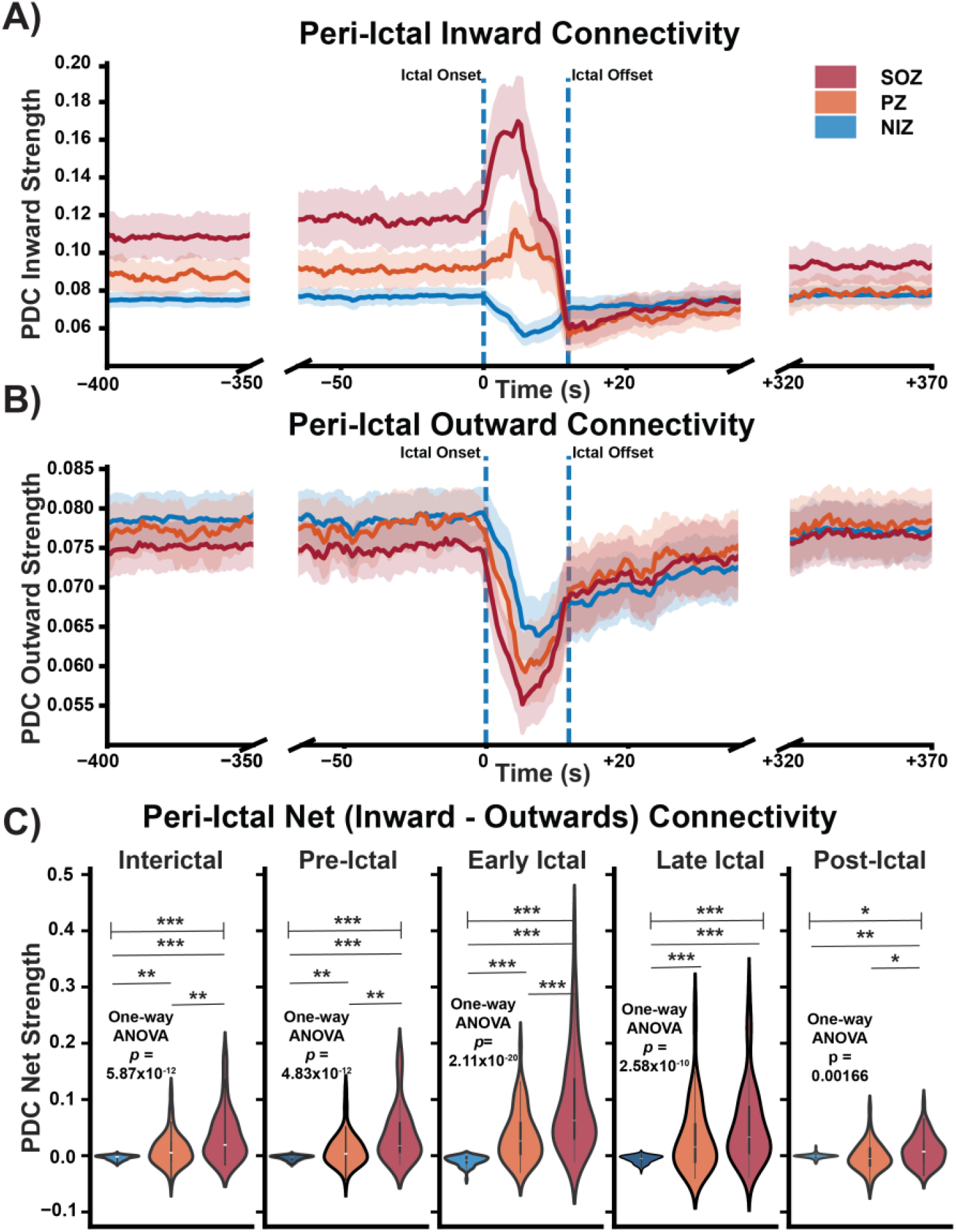
Directed connectivity demonstrates peri-ictal reorganization towards the SOZ and PZ. **A)** Inward connectivity is displayed across the peri-ictal period. **B)** Outward connectivity is displayed across the peri-ictal period. **C)** Net (inward – outward) connectivity for each period for each window is aggregated and displayed as summary distributions. SOZs are shown in red, PZs in orange, and NIZs in blue. Significance was assessed with a one-way ANOVA. Post-hoc pairwise comparisons were completed with two-sample t-tests with corrections for multiple comparisons. Significance for corrected cutoffs: *p<1.25×10^−2^, **p<1×10^−3^, ***p<1×10^−4^.

### Consciousness Sparing Seizures Exhibit Different Trajectories than Consciousness Impairing Seizures

Next, we sought to analyze if seizure type influenced the peri-ictal connectivity patterns noted. We found that when stratifying by seizure subtype, FIAS and FBTC seizure network trajectories more quickly transitioned into the second ictal phase, often occurring during the early ictal windows. FAS seizures still demonstrate a network collapse during phase 2 in the late ictal windows, however net SOZ connectivity decreases significantly earlier for FIAS and FBTC seizures compared to FAS. Seizure subtype trajectories can be significantly distinguished starting in the late-ictal window (**Fig. 3A**). We found that the SOZs exhibit different net connectivity between seizure types during the phase 2 of the ictal period (**Fig. 3A**, one-way ANOVA, p<0.01 after false discovery correction over the multiple windows). Specifically, FAS have greater net PDC than FIAS or FBTCS (**Fig. 3C**, p=0.04, p=3.0×10^−4^, two-sample t-test), and FIAS have greater net PDC than FBTCS (**Fig. 3C**, p=2.7×10^−4^). As seizures progress, consciousness-sparing seizures (FAS) can maintain higher directed connectivity into the SOZ compared with consciousness-impairing seizures (two-sample t-tests, p<0.05 after false discovery correction).

**Fig. 3:**
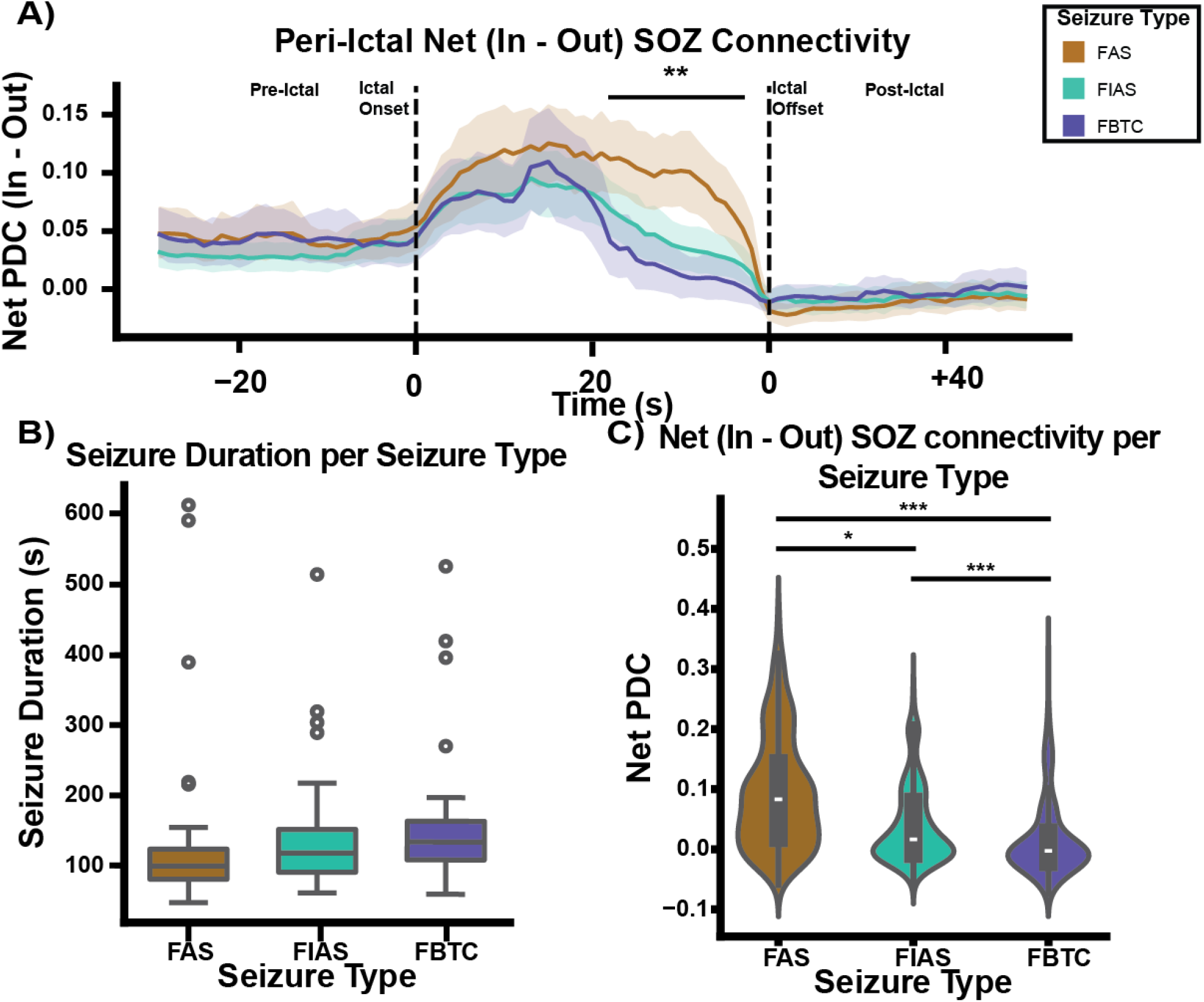
Loss of consciousness is associated with decreased ability to traffic communication to the SOZ peri-ictally. **A)** For all three seizure subtypes analyzed, net connectivity of the SOZ is plotted over the pre-ictal, early ictal, late ictal, and post-ictal period. **B)** Seizure duration in seconds is plotted for all three seizure subtypes. **C)** SOZ connectivity is aggregated and plotted as violin plots over the periods where consciousness-impairing seizures demonstrate significant differences between FAS compared to FIAS and FAS compared to FBTC (p=0.04, p=2×10^−4^, two sample t-test). *p<.05, **p<.01, ***p<.001.

Despite being aligned across start and end windows, it is possible that differences in seizure length may explain these network trajectories. Consciousness-impairing seizures (FIAS, FBTC) may last longer and thus have more opportunity to perturb and reorganize networks. Therefore, we quantified seizure duration and found that across our cohort we could observe no significant differences in seizure length across seizure subtype from one-way ANOVA (**Fig. 3B**). While this analysis does not associate loss of consciousness to a particular network profile or transition, we clearly observe that the peri-ictal network collapses sooner in seizures with impaired consciousness. This may suggest that seizure involvement is correlated with network collapse as consciousness-impairing seizures may be surrogate measures of broader network disruption.

### Seizure Spread and Directed NIZ Connectivity May Explain Seizure Phases

Despite the SOZ clearly exhibiting high net inward connectivity in phase 1, it is unclear which areas maintain this high communication. Although net SOZ connectivity diminishes in phase 2, it remains higher than net NIZ connectivity for the duration of the ictal period. At seizure offset, net SOZ connectivity is lower than all other connectivity **(Figure 2A-C**). Furthermore, these net increases in SOZ connectivity in phase 1 are not reflected by an increase in outward connectivity. The NIZ is the largest proportion of nodes sampled (**Table I**). When rebalancing against NIZ communication to itself, the PZ, and the SOZ, we observed that NIZ communication to the SOZ increased in phase 1 (**Fig. 4A)**. This implies that SOZ network dynamics were primarily driven by changes in NIZ connectivity.

**Figure 4.**
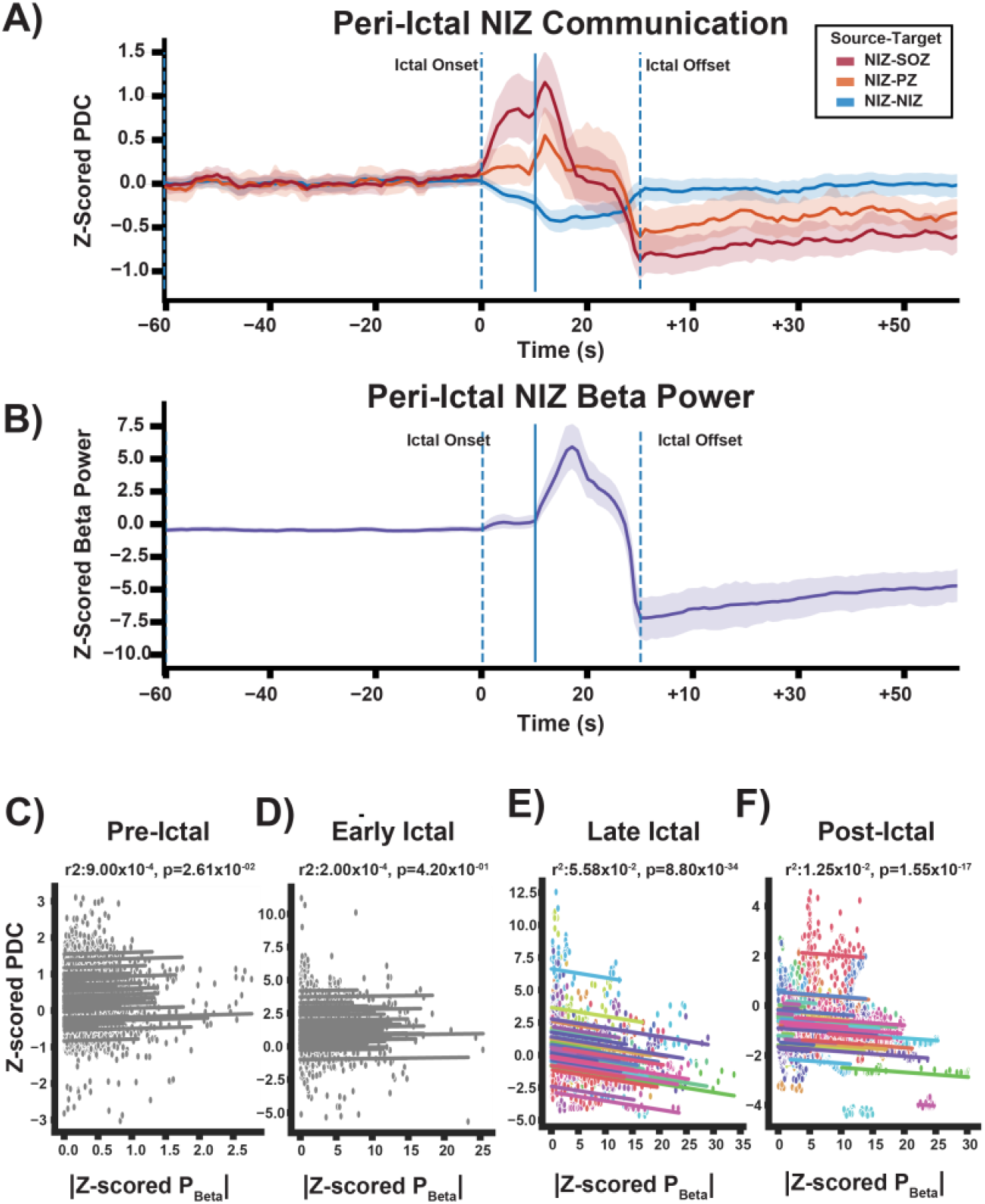
Seizure Propagation Correlates with NIZ Network Collapse. **A)** NIZ connectivity is highlighted with all channels normalized to zero for the interictal period. Connectivity from the NIZ to the SOZ is shown in red. Connectivity from the NIZ to the PZ is plotted in orange, and intra-NIZ connectivity is shown in blue. A vertical line indicates the time stamp where beta power begins to most rapidly spread across the NIZ. **B)** Z-scored beta power is plotted for the NIZ during the pre-ictal, ictal, and post-ictal periods. A vertical line marks the moment where beta power begins to most rapidly increase in the NIZ. **C)** Pre-ictal **D)** Early ictal **E)** Late ictal and **F)** Post-ictal correlations between instantaneous absolute beta power and instantaneous NIZ to SOZ connectivity is plotted. All subjects’ data are plotted as a scatter with a repeated measures correlation line fit over their respective data. Each subject is color coated. Pre-ictal and early ictal plots are greyed out because no significant relationships were observed.

### Phase 1 SOZ Inward Connectivity Increases as Intra-NIZ communication Decreases

Measuring average connectivity from the NIZ to the SOZ, PZ, and NIZ corrected for sampling imbalances between all 3 regions. Notably, we found that intra-NIZ connectivity decreases **(Fig. 4A)**. We also found that NIZ→SOZ connectivity increases in phase 1, mirroring the net connectivity findings in **Fig 2A-C**. Furthermore, NIZ→SOZ ictal dynamics were also mirrored by the PZ (**Fig. 4A**) but with a lower magnitude of directed flow from the NIZ. Given that NIZ regions greatly outnumber PZ and SOZ regions, a shift in connectivity between intra-NIZ connectivity to NIZ→SOZ and NIZ→PZ communication can increase the inwards connectivity of the PZ and SOZ while still appearing as a global decline in NIZ connectivity.

### Phase 2 Network Collapse Coincides with Seizure Spread

Tracing spreading beta power across the NIZ reveals an initial spike in beta activity in phase 1 followed by a plateau of activity (**Fig. 4B**). The inflection point of pathological beta propagation is marked as a solid, vertical line and coincides with the period where NIZ to SOZ communication is at peak before dissipating.

Z-scored directed flow from the NIZ to the SOZ and PZ begins decreasing in phase 2 (**Fig. 4A)**. Rapid increase in beta power coincides with this precipitous decrease in SOZ inward connectivity from the NIZ **(Fig 4A-B)**. Electrographic seizure offset is marked by another precipitous drop in NIZ beta power and a post-ictal state with drastically decreased beta power compared to interictal **(Fig. 4B)**. Repeated measures correlation regression reveals a net negative relationship between NIZ to SOZ connectivity and the absolute value of z-scored beta after seizure propagation (**Fig. 4E**). This observed negative correlation implies that seizure spread may be diminished by high network connectivity into the SOZ.

### Peri-Ictal Connectivity Data Can Classify the SOZ

After observing that the SOZ was most distinct from the NIZ and PZ in the interictal, pre-ictal, and early ictal phase (**Fig. 1C**), we utilized net channel-wise connectivity with data from each of these three periods to train an SVC classifier to determine which channels belonged to the SOZ. Training an SVC for patients with Engel I outcomes (N=23) then aggregating test set predictions yields an average AUC of 93% (**Fig. 5A**). Receiver operating characteristics are plotted with SOZ as the positive class and the NIZ grouped with the PZ as the negative class (**Fig. 5A**). Average performance remains above the chance level for all false positive rates.

**Fig. 5.**
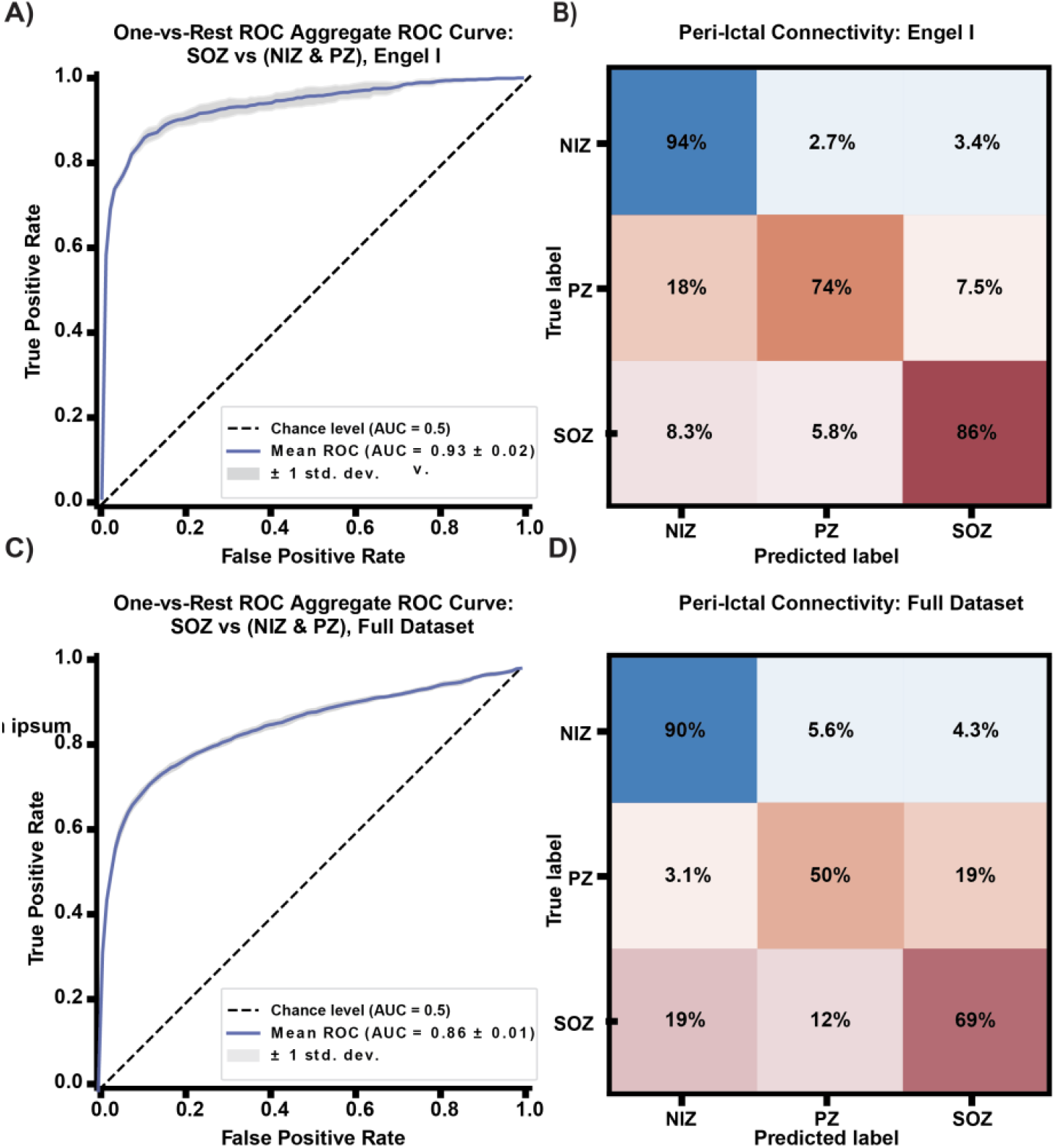
Peri-Ictal connectivity can localize the SOZ. **A)** For all patients with an Engel I outcome score a support vector classifier (SVC) was trained with nested cross validation and assessed on a held out test set. Aggregate SOZ vs NIZ and PZ receiver operating characteristics are plotted for all models. **B)** An average confusion matrix over all cross validation runs **C)** An ROC curve for the full dataset is shown **D)** An average confusion matrix over all folds of cross validation for all patients is shown.

Estimator variance is plotted as well and shows an average of 2% uncertainty. An average confusion matrix for each class shows high bias toward the NIZ with 94% accuracy for NIZ data in the test set. The PZ is classified with 74% accuracy. The SOZ is classified with 86% accuracy. For the full dataset we observed above chance classification for the SOZ with a combined AUC of 86% and SOZ classification accuracy of 69% (**Fig 5C, D**).

## Discussion

### All Seizure Types Exhibit Two Ictal Phases

In this investigation, we found that directed connectivity profiles for seizures follow stereotyped trajectories largely preserved across seizure types. These trajectories could be summarized as two distinct phases. In phase 1 we observed a coordinated increase in SOZ inward connectivity followed by phase 2 which presented primarily as a precipitous collapse of network connectivity to the SOZ. While consciousness-impairing seizures have differing timing for phase transitions, they presented overall similarities in organization between the NIZ and SOZ. Notably, consciousness-impairing seizures may be distinguished by their earlier network collapse. This earlier collapse in consciousness-impairing seizures suggests that large-scale network involvement drives the loss of inward connectivity to the SOZ.

### How Can SOZ Inward Connectivity Increase while Global Outward Connectivity Decreases?

While the SOZ experiences increases in communication during the early ictal period (phase 1), globally it appears that net communication drops. This finding implies that SOZs demonstrate diverging network trajectories from the global network in the ictal period. Nonetheless, at the summary level, we could not discern the source of increased SOZ connectivity. Further analysis suggests this increase is driven by NIZ-directed flow to the SOZ, while intra-NIZ connectivity diminishes. Within NIZ directed connections, intra-NIZ connections far outnumber NIZ to SOZ connections, thus a relative increase in NIZ to SOZ communication is masked when summarizing over all NIZ outward connections. High NIZ to SOZ connectivity aligns with a plateau in seizure spread. In phase 2, as seizures propagate to NIZs, SOZ connectivity deteriorates. The post-ictal collapse likely reflects a widespread decline in activity, with pathologic seizure spread disrupting normal communication patterns and impairing NIZs’ ability to sustain suppression of the SOZ.

### NIZ Network Abnormalities Precede Full Seizure Spread and Correlate with High Attempted SOZ Suppression

Analyzing connectivity phases against beta power highlights a possible causal relationship between inward connectivity to the SOZ and seizure activity. During phase 1, the SOZ is receiving its highest inward connectivity from the broader network. Relative to the interictal period, beta power within the broader network increases but plateaus during phase 1 period of high SOZ connectivity. This suggests that higher synchronous activity has perturbed the broader network and may have resulted in an increase in autoregulatory feedback. Nonetheless, as seizures persist this connectivity profile collapses and the seizure spreads. Thus while high inward connectivity may be a suppressive feedback mechanism to the SOZ, it is likely not sufficient for preventing all seizure activity. Furthermore, the network collapse indicates that broader network suppression is likely not responsible for the termination of seizure activity. The absence of a pre-ictal network perturbation indicates that seizure activity stems from SOZ-specific aberrancy rather than a change in inward connectivity. This challenges our prior postulate that a momentary lapse in the ISH-defined connectivity motif may create a permissive network state to allow seizure activity to spread. Instead, SOZs may independently transition into seizures. This finding has two implications for seizure forecasting studies. First, if seizure onset is a function of local activity, then these classical connectivity analyses are not sufficiently encoding node state. Second, networks may exist in high-risk states at larger timescales. Work analyzing chronic implants has shown that seizure risks fluctuate in a multidien cycle as do connectivity profiles.^33-35^ Thus, while there may be no pre-ictal change to network connectivity that permits seizure initiation, there may be a nuanced, long-term fluctuation in network profile which itself accommodates seizure initiation at any time within this high risk state.

### Augmenting the ISH with Peri-Ictal Dynamics

The findings in this investigation are concordant with our previous work on the ISH. In those investigations we surmised that high inward connectivity to the SOZ may be a consequence of network inhibition. Stimulating the broader network attenuated SOZ low frequency activity while augmenting high frequency activity. We interpreted electrographic attenuation to indicate suppression based on literature that highlighted GABAergic activity in this frequency band. In this study we find direct evidence of seizure propagation constrained by high SOZ inward connectivity. Prior to seizure spread to the broader network, the NIZs appear to focus connectivity to the SOZ. These observations augment the conclusions we drew from prior stimulation studies. Furthermore, it is possible that the outcome of this network suppression could be a critical determinant of the progression from focal aware to focal impaired aware seizures.

Specifically, increased SOZ connectivity appears to be maintained throughout the duration of FAS seizures while precipitously diminishing midway through consciousness-impairing seizures. This decrease in SOZ communication coincides with a continuous drop in intra-NIZ communication in phase 1 which precedes the zenith of seizure spread. Most NIZ nodes are sampling cortical regions. Therefore, the ability to suppress the SOZ by the NIZ may also be a parallel mechanism to explain impairment of consciousness. Prior work on the network inhibition hypothesis (NIH) has also shown that seizure spread to crucial arousal structures may explain loss of consciousness for patients with FIAS.^36, 37^ In this manuscript, we investigate another possible source of cortical: insufficient inhibition of the SOZ. While the NIH explains impairments in FIAS, the consequent drop in intra-NIZ communication is seen in FAS and FBTCS. Thus, these findings may offer a complementary mechanism to explain cortical impairment across seizure types (**Fig. 6**).

**Fig 6.**
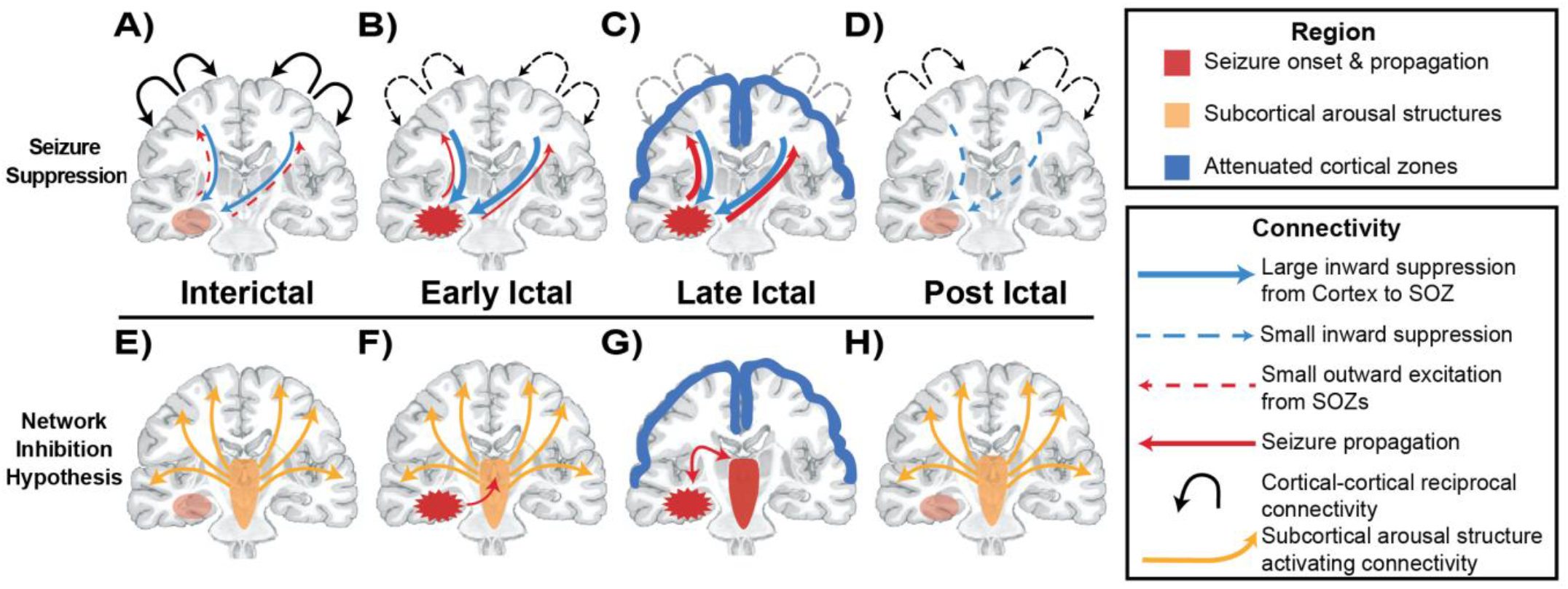
The Seizure Suppression Hypothesis in focal epilepsy. **A)** Network inhibition extends into the ictal period and may explain some of the aberrancy during seizures. **B)** Specifically, at seizure initiation, the broader network responds to overexcitation and hypersynchrony by trafficking higher inhibitory signal back into the SOZ. **C)** As seizures progress, network suppression is overwhelmed and normal intracortical communication diminishes which also may coincide with loss of consciousness. **D)** In the post-ictal state, the non-involved network slowly recovers but has not immediately reached pre-ictal inward connectivity to the SOZ or intracortical communication. The Network inhibition Hypothesis (NIH) **E)** seeks to explain why focal seizures may lead to loss of consciousness. In the interictal state, the brainstem sends tonic activating signals to the cortex to maintain consciousness **F)** As seizures initiate, they spread to brainstem structures **G)** and mitigate its ability to maintain arousal signaling thus leading to loss of consciousness as the cortex adopts a default delta rhythm. **H)** As seizures terminate, the ascending arousal structures in the brain stem are released to resume sending tonic activation to the cortex.

### Peri-Ictal Connectivity Distinguishes the SOZ for Localization

In prior work, an SVC model trained on interictal connectivity achieved an AUC of 84% for all patients. Adding peri-ictal data improved model performance to 86% with a more balanced classification schema, despite the highly imbalanced data (often 2 SOZ channels per 100 NIZ channels). For Engel I outcomes, the classifier achieved higher overall accuracy (AUC = 93%) and more balanced NIZ performance (94% vs. 65%) compared to previous work.^20^ These models were tasked to learn seizure specific SOZ designations and not the final SOZ designations, which makes the classification more difficult. However, this training and validation configuration was designed to simulate the information at hand for the clinical team during an EMU stay. We expect that the ecological validity of this configuration will make these findings most readily adapted to the clinical setting.

### Revisiting the Role of the Non-Involved Zone

Similar to prior work, we find that the SOZ and the NIZ appear to engage antagonistically during the ictal period.^38, 39^ In prior work by *Khambhati et al*., seizure propagation was studied as a function of nodes that drove network synchronization.^34^ They observed that seizure spread inversely correlated with a network’s ability to maintain diffuse connections. Interestingly, non-SOZ nodes exhibited the most variation in (de)-synchronizability, and thus were likely allowing seizure spread through high synchronization or low seizure spread by maintaining desynchronization within the network. The present investigation corroborates this push-pull dynamic through windowed network analysis. We also highlight the broader network’s in seizure suppression. Furthermore, aligning network dynamics across all 75 patient’s seizure events allowed us to discern specific phases of this push-pull relationship.

Conventional SOZ designation is guided primarily by the earliest recordings of seizure activity. Nodes which initiate seizure activity are designated SOZs, whereas nodes that pick up pathological activity within the first 10s of seizure activity become PZs. By default, the rest of the network is deemed the NIZ. This designation of NIZ, while apt in highlighting the areas which do not initiate seizures, may be due for refining. While SOZs begin seizures, evidence is mounting that the NIZs permit them to spread. There are likely nodes with different susceptibilities to pathological recruitment. This antagonism between directed connectivity and seizure spread calls for future investigations to differentiate types of NIZ nodes and possibly for a new designation within the NIZ. These findings also offer a starting point for neuromodulation. Nodes at the extremes of susceptibility/robustness may be viable targets for neuromodulation. Perhaps some nodes in the NIZ may be redefined as robust nodes if they consistently resist seizure spread. Susceptible nodes may be defined as those which most consistently succumb to seizure activity. If a paradigm could bolster a susceptible node’s ability to resist seizure spread or a robust node’s suppressive inward connectivity to the SOZ, then clinicians may be able to limit seizure spread and thus impairment.

### Extending the ISH to Explain Ictal Cortical Dysfunction in Drug Resistant Epilepsy

Prior work has found that loss of consciousness may be maintained by excessive synchronization between cortical and thalamic structures after propagation to subcortical arousal structures.^24, 37^ While our findings elucidate diminishing intra-NIZ communication, this difference may be reconciled by the construction of connectivity. Since PDC accounts only for non-zero phase lag in signals, it is possible that synchrony is discounted in the construction of PDC based connectivity. This difference may also be explained by thalamocortical synchronization,^24, 40^ which we are not powered to analyze within this cohort. Additionally, diminishing intracortical communication is not mutually exclusive to heightened thalamocortical coupling, and may in fact be a corollary finding that explains loss of conscioussness.^41^

The NIH suggests that disruptions in connectivity from brainstem to cortical structures underlie consciousness impairments. We propose a complementary mechanism: failure to sustain inward suppression to the SOZ. (**Fig 6**.) Our findings extend the ISH to a broader seizure suppression network active beyond the interictal period, where cortical responses aim to inhibit the SOZ and counter seizure initiation. Consciousness impairment may result from insufficient NIZ-directed suppression, allowing seizure propagation to subcortical arousal structures. Prior studies examining GABAergic tone in seizure onset note that seizure initiation is marked by an attempted rise in inhibition. Our work supports this finding and highlights the crucial role network inhibition may play in combatting seizure propagation.^22^

### Limitations

Important limitations to this work mainly involve the signal analysis assumptions made to analyze these data at the group level. Specifically, aligning all seizures across time requires some summarization and it’s possible that mid-window contractions have left out some crucial network transitions. We attempted to account for this contraction by generating distributions with balanced resampling over large windows for all periods, however this view does not allow for clear trajectory comparison. When examining seizure subtypes, we are not privy to the exact moment that consciousness becomes impaired, thus we are unable to determine if this transition in clinical symptomology correlates with a clear network transition. Future work with data labelled at this time point may focus on aligning network states to this moment as a zero point, instead of seizure onset. Classifying nodes as SOZ vs non-SOZ creates a highly imbalanced learning problem. While nested k-fold cross validation limits overfitting, class imbalance will persist. This analysis was performed on a single institution’s data. Multi-site validation can offer a control against systematic biases in data collection, storage and pre-processing.

## Conclusions

Directed connectivity analyses suggest that the ictal period is marked by 2 distinct phases: 1) a diversion of communications into the SOZ with a coincident decrease of normal broad network communication followed by 2) a second phase where all network communication collapses, including inward to the SOZ. Seizures may evolve to impair consciousness from failure to successfully suppress ictal spread. Prior work on the ISH suggests that the NIZ actively directs communication into the SOZ in an inhibitory manner. This work extends the ISH into peri-ictal network dynamics. By correlating beta-scored seizure involvement with network connectivity, we observe a possible antagonism between directed connectivity and seizure spread. It is possible that seizure involvement is mitigated initially by a diversion of resources towards network suppression. Intracortical impairment may be the result of insufficient seizure suppression. Further work is needed to examine this possible causal mechanism of seizure suppression. Perhaps this evidence may redirect the aim of neuromodulation studies to apply stimulation to the broader network to bolster innate seizure suppressive networks. Overall, this work is in line with current network studies which marks a clear departure from focusing solely on the SOZ.

Finally, we contend that this work makes the case for enriching how we describe these broader seizure networks, as their peri-ictal connectivity profiles clearly demonstrate an “involved” role in seizure propagation.

## Acknowledgements

A complete draft of the manuscript was reviewed by ChatGPT-4o, and after careful review, some edits were adopted to improve grammar and word choice.

